# Temporal Characteristics of Neonatal Chick Retinal Ganglion Cell Responses: Effects of Luminance, Contrast, and Color

**DOI:** 10.1101/2022.07.28.499133

**Authors:** Deepak CS, Abhijith Krishnan, K S Narayan

## Abstract

We perform microelectrode array recordings of neonatal chick retina explants and report stimulus-dependent response properties of different retinal ganglion cell (RGC) types to various luminance and contrast adaptation conditions. The isolated single units indicate wavelength-sensitive response properties for all the retinal ganglion cell types recorded from. Responses to different luminance and contrast conditions investigated as a function of wavelength indicates a combination of adaptation and sensitization features in the recorded population of RGCs. We further demonstrate the presence of complementary response properties in most of the RGCs to blue (450 nm) and green (530 nm) light input and infer that luminance, contrast and color information are encoded in a wide variety of metrics such as latency, response event duration and pairwise correlations.

## Introduction

Gallus gallus domesticus represents a diurnal vertebrate with a cone-rich retina consisting of an afoveate area centralis with a high density of single and double cones and a periphery with double cones and rods. (Seifert et al., 2020) The chick-retina has been described as a model avian retina and has been recognized as a suitable cone rich retina to study eye disease and development. (Wisely et al., 2017). The importance of studying the chick retina has been highlighted in recent reviews to get a generalized understanding of the working of the vertebrate retina. (Baden et al., 2021) The structurally dense chick-retina features many neurons with complex branching patterns that divide both outer and inner plexiform layers into multiple anatomical strata, which presumably have distinct functions. The chicken follows the typical vertebrate retinal organization, with three nuclear layers: the ganglion cell layer (GCL), inner nuclear layer (INL), and outer nuclear layer (ONL), and three neuropils: the retinal nerve fiber layer (NFL), inner plexiform layer (IPL) and outer plexiform layer (OPL). Signal processing occurs along two major pathways. In the vertical pathway, bipolar cells connect photoreceptors to the retinal ganglion cells whose axons form the optic nerve. Horizontal pathways formed by horizontal cells in the OPL and amacrine cells in the IPL, modulate the vertical pathway, mediating processes such as lateral inhibition and directional selectivity. The high density of RGCs in the area centralis and the dorsal area with cone to RGC ratios averaging 1.5, resembles the primate parafovea providing high visual acuity. (Seifert et al., 2020) The understanding of chicken retina development is beneficial in the context of prosthetics that elicit specific activation of different RGC types. (Deepak et al., 2022)

Though there is now a recent wealth of anatomical information available on the chick retina with recent studies done on its connectivity in the outer plexiform layer (Gunther et al., 2021) and solving of the chick transcriptome (Yamagata et al., 2020), there is a scarcity of electrophysiological studies performed on it. The utility of multi-electrode array extracellular recordings on neonatal chick retinal explants to extract the wavelength-dependent response properties of different RGC types is presented. The effect of the stimulus protocol on the elicited responses for various putative RGCs is described with responses showing dependence on ambient luminance, pulse width, and the duty cycle of the periodic full-field flash.

## Results

Retinal explants were probed with different full field stimulus protocols to extract different temporal characteristics of chick retinal ganglion cell responses. Explants were presented with the following stimulus protocols:

1. White flashes across luminance levels and different duty cycles.
2. Colored full field flashes of different pulse widths.
3. Constant illumination at appropriate ambient light intensities to gauge color dependent luminance adaptation effects.
4. Random Gaussian full field flicker stimuli with the mean corresponding to the ambient illuminance and the variance of the distribution corresponding to the contrast, to extract the spike triggered average (STA) which represents the linear response of the putative RGC recorded from.
5. Alternating random Gaussian flicker stimuli with the same illuminance and different contrasts to probe color dependent contrast adaptation effects.

The chick retina has been reported to describe spatial inhomogeneities in the RGC mosaic. (Baden et al., 2020) To reduce variability across different preparations, retinal explants were sampled from the peripheral dorsal area across twenty explant preparations, from which 969 single units were isolated through spike sorting and clustering using SpyKing circus (Yger et al., 2015). There are intrinsic difficulties of recording from the chick retina owing to waves of spreading depression[] after the onset of which the fundamental firing properties of RGCs change and the native response state is nearly impossible to recover. In our measurements only samples that last over 45 minutes before the onset of the spreading depression event are considered for the analysis.

### Response Polarity

The response of chick retinal explants to full-field flashes and random flicker stimuli yielded a range of response types that differed in transiency, latency, and trial-to-trial variability. Full-field flash stimuli at different luminance levels from the dark were provided to test the responsive nature of the retinal explant. Initially, responses to white light for intensity varying in the range of 0.7 to 70 mW/m^2^ are probed.

In this simplified scheme, RGCs are classified functionally based on their response polarity, i.e., they are ON RGCs if they fire to increasing contrasts, OFF RGCs if they fire to decreasing contrasts, and ON-OFF cells if they fire to both. **Figure 1** illustrates different polarity of response observed in the different types: ON, OFF, and ON-OFF. **Figure 1(a-c)** shows that the response polarity is clearly demonstrated upon having both positive and negative steps around a photopic mean light intensity. Responses predominantly peaked at the switch from high to ambient luminance but showed little to no activity when the luminance switched from ambient to dark. This is custom pulse shows better response in comparison a simple positive contrast pulse observed over short time scales (Figure S1). However, it is to be noted that a positive contrast would provide a better indication of the response type when measurement over long time scales. **Figure S2** shows that the response polarity is clearly discerned for “1 s ON – 5 s OFF” measurement scheme. This flash scheme is considered for all the subsequent measurements presented in the paper.

**Figure 1.**
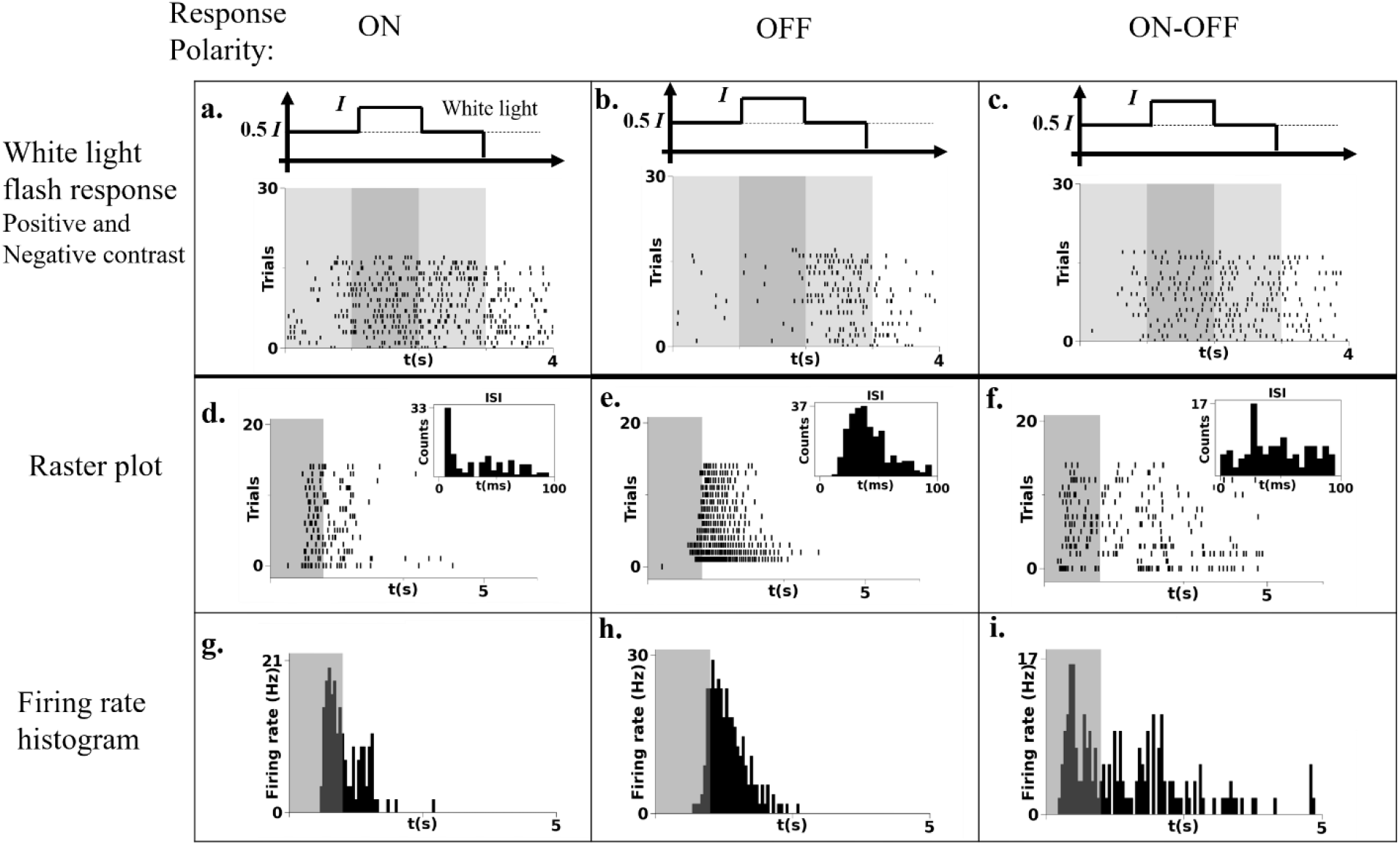
The response of 3 representative retinal ganglion cells to 20 trials of sequential positive and negative 100% full field contrast steps of a white LED flash around a photopic mean luminance of 7 mW/m2 .(a)Raster plot of a predominantly ON type cell,(b)an OFF type cell and (c)an ON-OFF cell type are described. The raster plot of a representative (d)ON type, (e)OFF type and (f) ON-OFF type responses to ten trials of a 1 s ON – 5 s OFF (20% duty cycle) white LED flash stimulus.(g-i) The firing rate histogram plots of the raster plots described above in (d-f)

**Figures 1(d-f)** and **Figure 1 (g-i)** shows the raster plot and the corresponding firing rate histogram (FRH), respectively, for the representative RGC types for the ON, OFF and ON-OFF polarities. The long OFF time is a suitable method since it provides a longer time for the neuron to reset to the native state, thereby allowing for lower variability across trails. In our studies over numerous explants, a large percentage of RGCs extracted (>90%) in the neonatal chick retina described a slow and sustained component in the response at some intensity level.

In addition to broadband white light responses, the polarity based firing response were also observed upon having monochromatic illumination. **Figure S3** shows the representative response to blue light (430 nm). Broadly, the polarity-based classification of the RGC types holds consistent. It is also important to note that some cell types showed a change in response polarity with changing ambient illumination levels (**Cell 2 Figure S4**) while some others retained their response polarity. (**Cell1&3 Figure S4**) (Tikidji-Hamburyan et al., 2015)

### Response Kinetics

In addition to the polarity of the firing responses, the temporal kinetics is a useful parameter to further classify the RGC response. **Figure 2** illustrates the dominant kinetics associated with the different RGCs reported. Here we represent the different kinetics corresponding to two different response polarity: ON and OFF. **Figure 2a** and **2b** shows a fast and delayed ON response to the blue full field flash, respectively. **Further, 2a** shows transient (phasic) decay features in the firing rates, while **Figure 2b** shows tonic features in the firing rates. **Figure 2c** shows a mixture of the phasic and tonic features in the response. This is evident in the distribution of the inter-spike interval (ISI) shown in the figure insets. Similarly, **Figure 2(d-f)** describes the phasic, tonic and mixed kinetics observed in the OFF-type RGCs.

**Figure 2.**
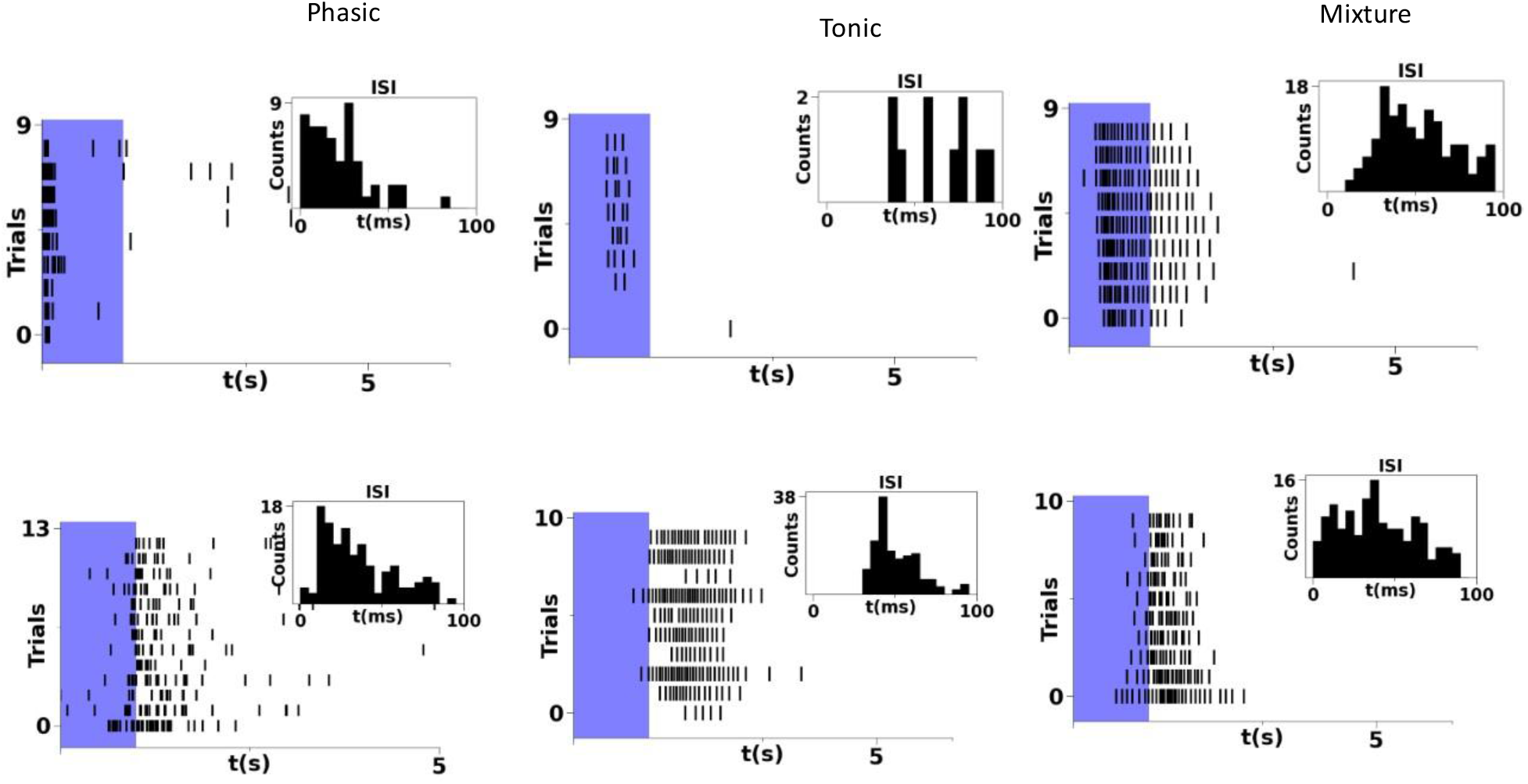
Tonic and Phasic response types(a)The response fast ON cell showing phasic firing features in response to 9 trials of a 1s ON-5 s OFF blue LED light flash. The low inter spike interval (ISI) that is characteristic of phasic firing described in the inset. (b) The response of a slow ON cell type describing tonic firing features to the same 20% duty cycle blue LED flash stimuli.(c) The response of a sustained-ON ganglion cell type to blue light flashes describing a mixture of phasic and tonic firing features, the ISI histogram shown in the inset indicates a broad distribution.(d) Raster plot of an OFF-type ganglion cell describing phasic firing properties to blue LED flashes. (f) The response of a tonic firing OFF ganglion cell to the same stimuli. (f) The response of a predominantly OFF ganglion cell showing a mixture of tonic and phasic firing features.

Intensity dependence of the response for a white light stimulus is shown in Figure S4. A corresponding overall increase in average firing across different single units is observed as the white stimulus intensity is increased. Yet, the firing rate shows an opposite trend in certain cell types (**Figure S4c,f**).

To further observe the temporal kinetics spike firing statistics on the pulse width was probed for different RGC types. Explants from the dorsal area of the neonatal retina with full field flashes of varying pulse widths was probed and the activity is monitored for 5 s. **Figure 3(a) and (b)** shows the raster plot of a given cell for a green flash with pulse duration of 50 ms and 500 ms, respectively. The ON-transient nature of the response is independent of the pulse duration. In contrast, Figure 3(c) and 3(d) shows pulse duration dependent responses. In addition to the transient component, a pulse-width dependent sustained component is evident. The firing rate histogram shown in **figure 3d** describes characteristic bimodal ON-OFF responses to green.

**Figure 3.**
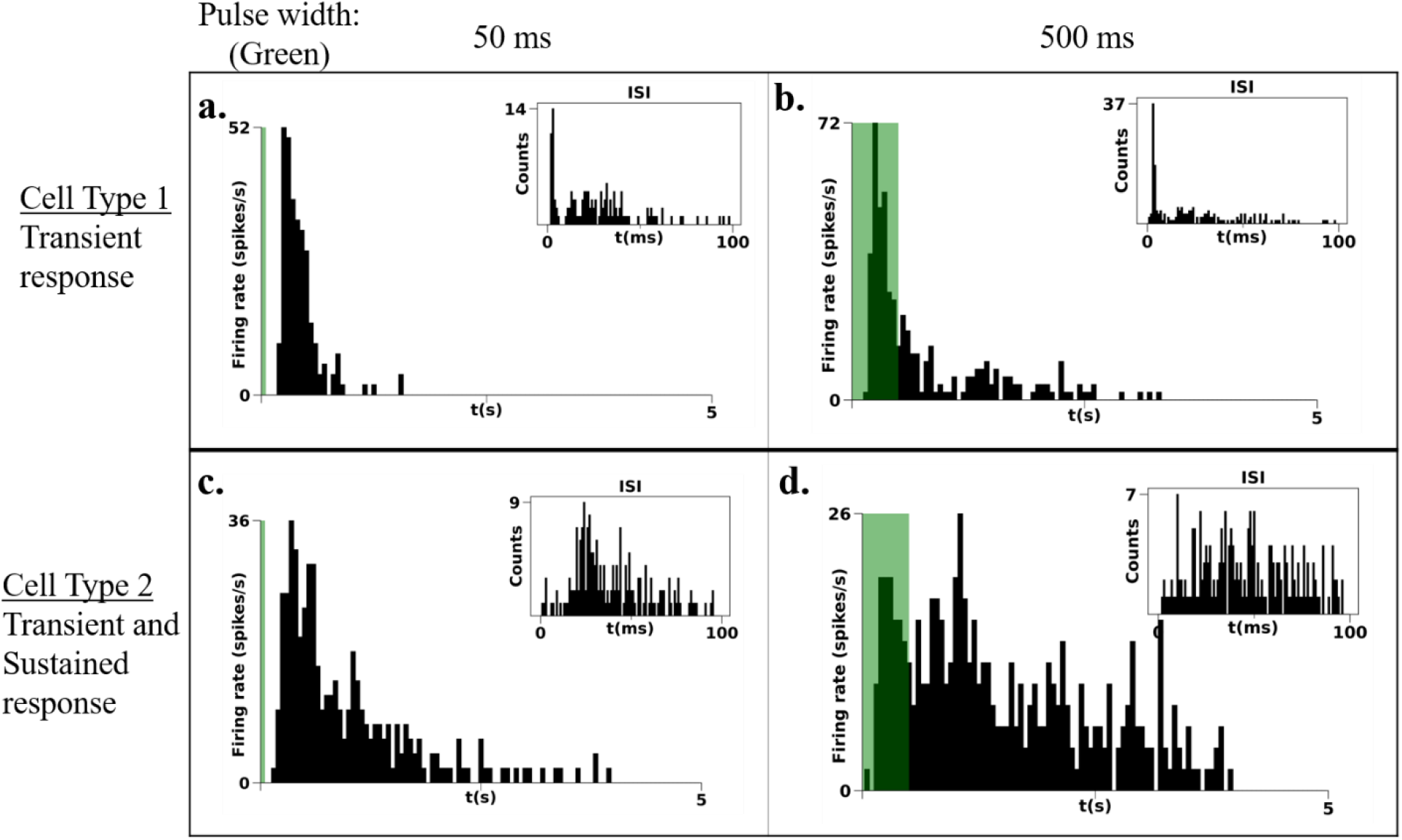
The response of a transient ON RGC to (a) 50 ms-ON 5s-OFF green LED flash and (b) the response of the same ganglion cell to 500ms periodic green LED flash with a duty cycle of 5 seconds. The response of the ON transient is invariant of the pulse width of the full field flash. The response of a representative OFF ganglion cell to a (c) 50 ms flash with a duty cycle of 5 s and the response of the same ganglion cell to a (d) 500 ms light pulse delivered periodically with a duty cycle of 5 s. The OFF RGC shows qualitative changes in firing in the anti-preferred contrast with varying pulse widths. The inset in (a-d) is the ISI histograms.

A large number of delayed ON responses were found across explants samples and were observed for contrast steps of all wavelengths. For example, **Figure 4** describes the dominantly observed delayed ON- fast OFF response. Here the ON component depends on the pulse duration, while the fast-OFF component is observed to be pulse duration independent.

**Figure 4.**
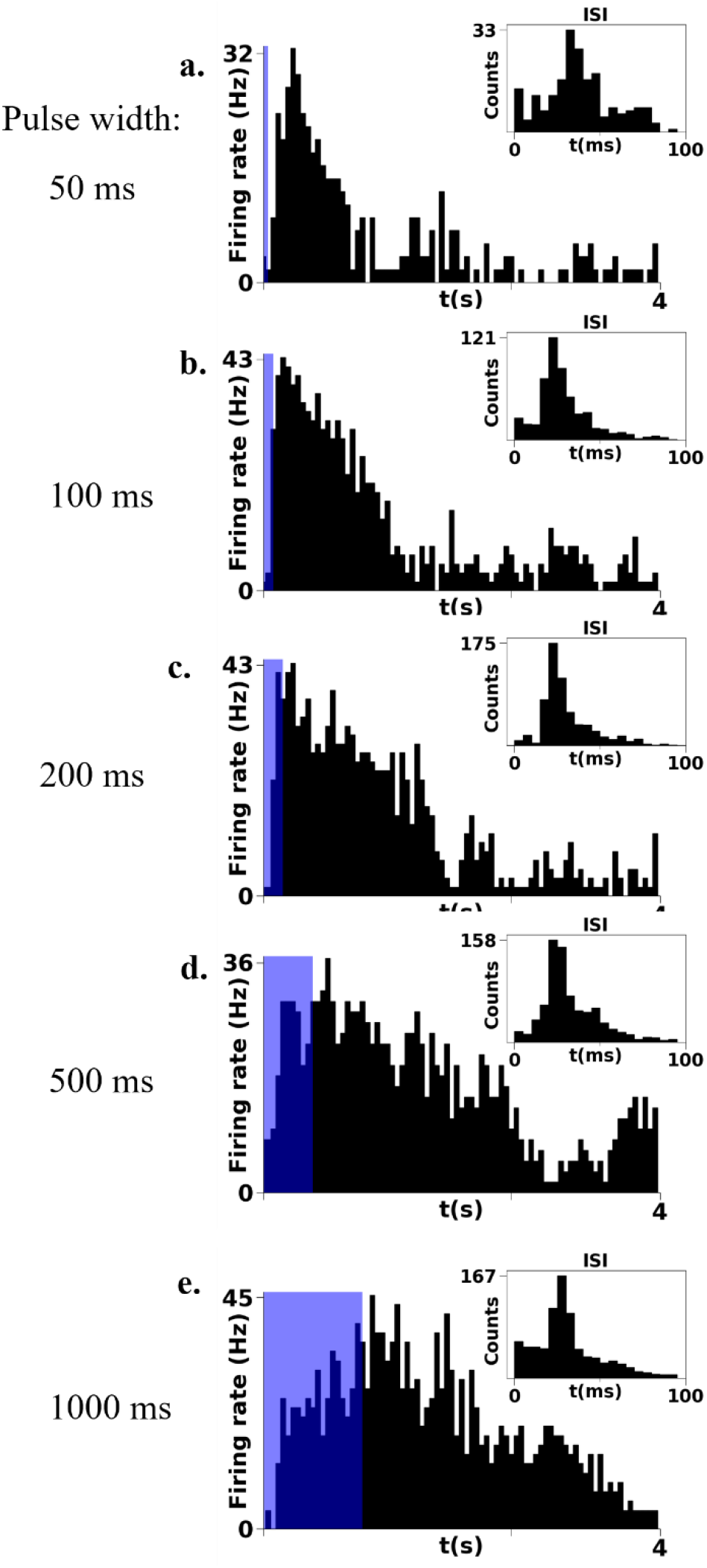
The response of a representative retinal ganglion cell describing both pulse width dependent sustained response and a pulse width independent transient response described for blue led flashes at an intensity of 1mW/m2 for different pulse widths (a)50ms,(b)100ms,(c)200ms, (d)500ms and (e)1000 ms with a fixed long OFF cycle duration of 5 s. The insets describe the ISI histogram for the respective flash responses across the 10 trials for the different LED flash pulse widths.

These were the most numerous and diverse response type observed with the firing rate decay profile of the ON and OFF components varying with wavelength, luminance and the duty cycle of the periodic flash.

### Adaptation and Sensitization

Adaptation and sensitization are key features of sensory systems to optimize the firing response required to reliably encode the statistics of the sensory input (Shapley et al. 2009), allowing the retina to respond to a large range of luminance levels. Adaptation and sensitization are mechanisms utilized by sensory systems to enable the dynamic modulation of the gain of different populations of RGCs by using different forms of short-term plasticity. This is achieved by the differential depression and facilitation of different excitatory and inhibitory synapses of bipolar cells in the retina. (Nikolaev et al., 2013)

Higher luminance values elicited a higher peak firing rate in all RGCs recorded and the same contrast changes at higher luminance levels also doing the same, pointing to the increased contrast sensitivity at higher ambient luminance values. (Chen et al. 2005) The adaptation and sensitization time scales varied for different cell types and also depended on the wavelength of the stimulus. To probe the Luminance Adaptation processes of different cell types, we recorded the firing activity to a steady state full-field illumination. **Figure 5(a-c)** shows the representative types of adaptation in response to steady state green light illumination. The observations indicate dominantly the two basic features.

i. An initial peak firing following a steep reduction over a period of a few seconds can be attributed to adaptation 5(a)) and was observed in a significant number of cells.
ii. Many cells also describe a *sensitization* feature (Figure 5(b)), which shows a delayed increase in firing rate occurring after the onset of the illumination.

**Figure 5.**
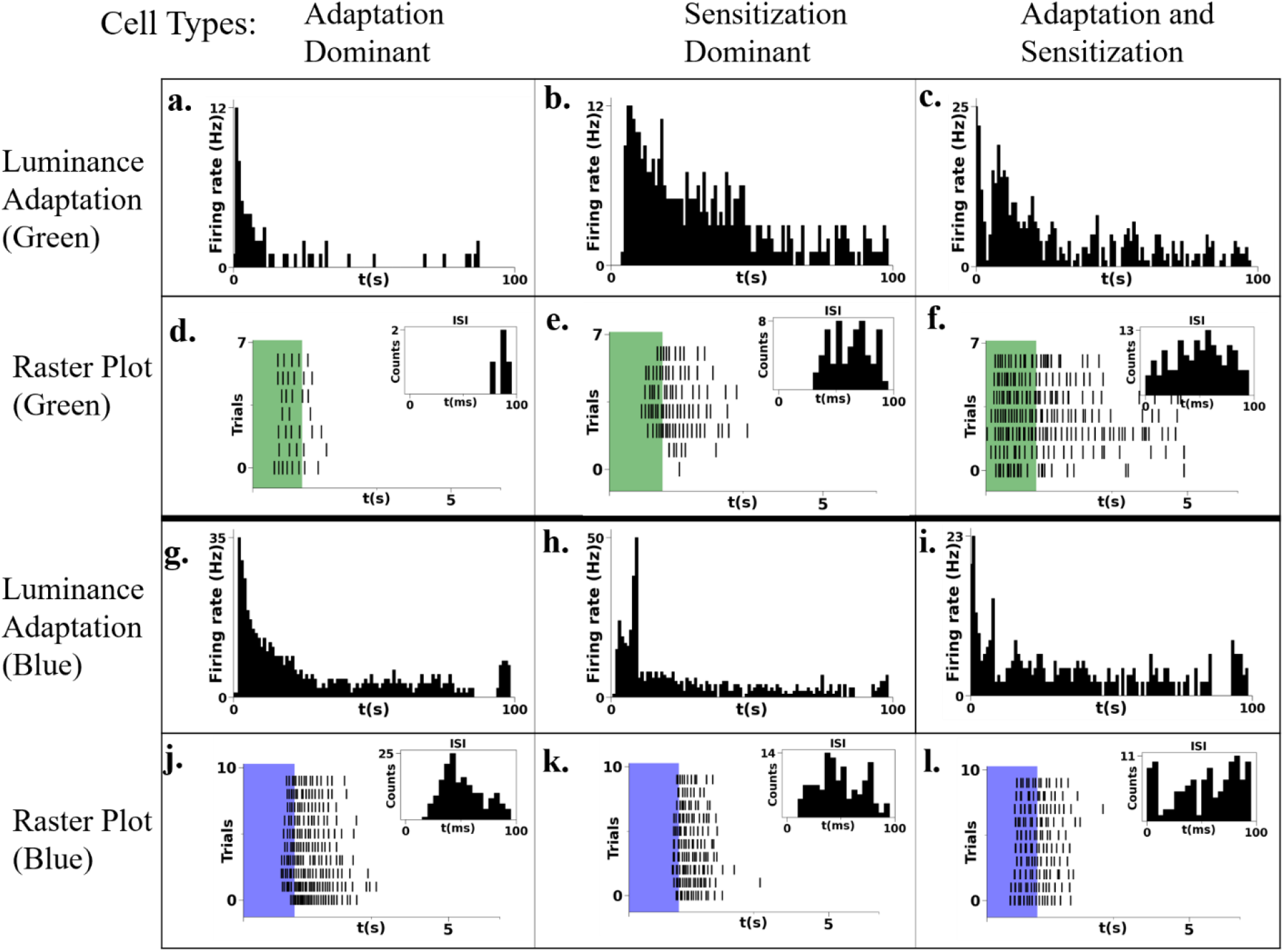
Luminance adaptation to uniform blue and green light illumination. (a) The firing rate histogram of a representative RGC type showing a sudden decrease in sensitivity and thus a rapid firing rate reduction thus describing slow luminance adaptation to uniform green LED illumination. (b)A representative RGC showing a delayed increase in the firing rate after exposure to uniform green LED illumination describing the sensitization feature. (c) A representative RGC describing both the slow luminance adaptation and the sensitization features. (d-f) The raster plots of the RGCs to full field 100% contrast steps of the adapting, sensitizing and both adapting and sensitizing RGCs described above. The firing rate histograms of representative (g)adapting, (h)sensitizing and (i)both adapting and sensitizing RGC types to uniform blue LED illumination.(j-l)The raster plots describing the full field flash responses of the representative RGCs described above in (g-i) to blue LED light.

Many of the cells indicate a combination of the above mentioned features and is represented in **Figure 5c.** Figures 5(g-i) describes the luminance adaptation plots for the representative cells to steady state blue illumination. It is noted that sensitization features are dominantly observed in the same cells upon have blue illumination. A consistency of the kinetics is evident with the corresponding pulse based raster plots (**Figures 5(d-e)** for green and **Figures 5(j-l)** blue illumination, respectively). In addition to ON-type polarity described in **Figure 5**, **Figure S5** shows the adaptation and sensitization features for an OFF-type RGC.

In addition to luminance adaption, a more realistic parameter to describe the adaptation kinetics is in response to light contrasts. Here, we study the contrast adaptation, i.e., temporal response of the RGCs to light intensity that dynamically varies around a mean intensity. The intensity flicker is governed by Gaussian white noise statistics. **Figure 6** describes the luminance adaption and contrast adaptation of the representative RGCs types classified on the basis on dominant adaptation and sensitization features, in response to green light stimulus. **Figure 6b**, **6d** and **6f** show contrast adaptation, sensitization and combination of both features, respectively.

**Figure 6.**
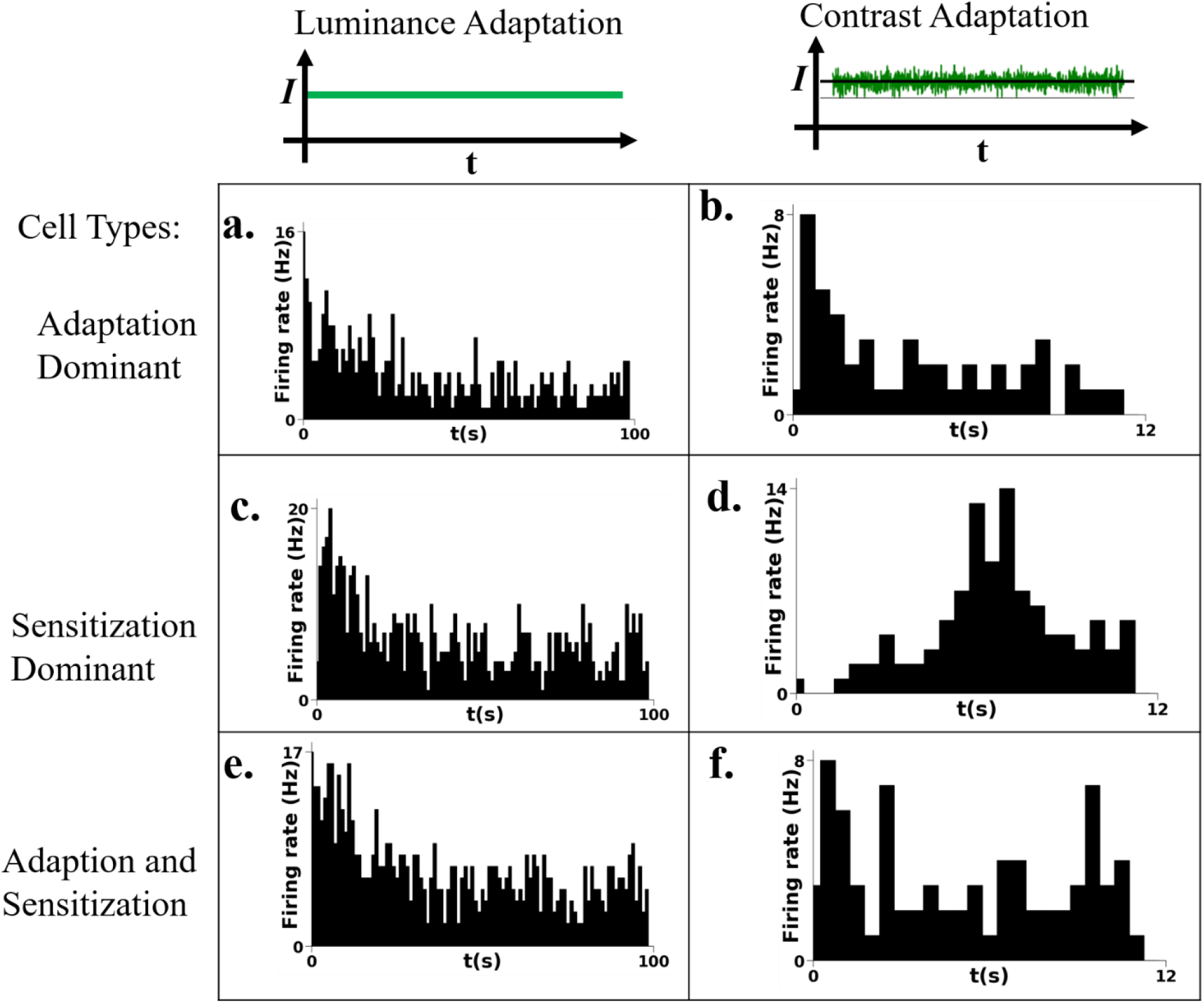
The firing rate histogram of an adapting RGC to (a)uniform green light illumination and (b)the response of the same RGC to presentation of high contrast green gaussian flicker stimuli(50% variance) averaged across 10 different gaussian sequences describing slow contrast adaptation. (c) The response of an RGC showing sensitization to uniform illumination,(d)and the response of the same RGC to high contrast gaussian stimuli describing sensitization to high contrast i.e adaptation of inhibition. (e,f) The response of a representative RGC describing both adaptation and sensitization features to uniform illumination and high contrast gaussian stimuli respectively.

Green light illumination predominantly resulted in only a fast adaptation decay of the order of a few seconds in only ON cells while it showed a slow/broader sensitization rise feature that ensues after the adaptation decay in ON-OFF cells. Blue and violet light resulted in sharp adaptation and sensitization features in the response to constant illumination, but these features were largely absent when the retina was exposed to high contrast, pointing to the fact that blue and violet light responses were more likely mediated by sustained bipolar cells and had a larger integration time prior to firing. The sensitization feature generally showed a peak between 3- 20 s depending on the wavelength and the cell type, with blue and violet cells generally describing a faster sensitization decay profile than adaptation decay and white and green responses showing a slower sensitization feature and a sharp adaptation feature. All neurons showed one or more of these features, with slow OFF cells showing only the sensitization feature, fast ON cells showing only the luminance adaptation feature and all other RGCs showing a linear combination both features.

Slow adaptation and sensitization both show clear changes in the firing rate with changing contrast values. (Kastner et al. 2013) The time scales of the slow contrast adaptation and the onset of sensitization seem to show significant overlap in the population of cell types that describe both features. This is indicative of both these complementary phenomena working to increase the dynamic range of operation of the retina as a whole; with sensitization compensating for the loss of sensitivity after a high contrast switch due to contrast adaptation and adaptation compensating for the saturation of responses due to sensitization.

Luminance and contrast adaptation has been studied extensively and modeled to understand the underlying functionality. Contrast adaptation in particular has been studied for fast OFF cells in many vertebrate retinae (Baccus and Meister 2005) and seem to show a slow and fast adaptation timescale. Intracellular recordings of the rod bipolar- AII amacrine cell synapse have shown that a common vesicular transport mechanism is shared by luminance and slow contrast adaptation. (Jarsky et al 2011) In addition the adaptation to the mean light level also involves Ca^2+^ channel inactivation mechanism.

### Response to Gaussian White Noise

Subsequent to the response studies of the full field flash input, the response to the random gaussian flicker stimulus was studied to gauge the repeatability and temporal jitter in the elicited firing events. Most of the clustered RGCs were responsive to both increments and decrements in contrasts and, if classified based on the R-ON/ R-OFF (bias index) metric, can be labeled as ON-OFF cells. (Deepak et al. 2022) This bias index metric was also observed to be highly sensitive to ambient illumination (luminance) and pulse width of the full field flash, indicative of integrated processes of the photons/charges generated in the RGC receptive field. More rigorous characterization of cell types was carried out by introducing a full field random flicker stimulus where intensity values of the LED were picked from a gaussian distribution with an effective mean luminance intensity of 1, 5 mW/m^2^ for the blue, and green LEDs respectively and a contrast factor controlled by the variance of the distribution. The linear component of the RGC response was then estimated by performing a reverse correlation of the random flicker stimulus with the elicited spiking events of the putative RGC. The spiked triggered average (STA), the average temporal contrast profile of the input that triggers a reliable RGC firing and has been used extensively to characterize RGC types across development. (Hilgen et al., 2017) Accordingly, we classify the polarity of the RGCs based on the most recent polarity of the spike-triggered average (STA) estimated from a Gaussian full-field flicker protocol.

**Figure 7** describes the firing response properties of a representative ON and ON-OFF RGC to green stimulus. In the case of a ON-type RGC**, Figure 7(a)** shows a fast ON response in the raster plot**. Figure 7b** shows the Post-Stimulus Time Histogram (PSTH) of the same RGC to ten trails of the same random GWN sequence (50 ms refresh rate, 1200 frames). The STA shown in Figure 7c indicates ON contrast polarity as evidenced by the positive peak prior to spiking (at t = 0). Ten trials (**Figure 7 (b)**) revealed the reproducible post stimulus time histogram (PSTH) of the same fast ON cells. This method of analysis holds for ON-OFF cell types as described in **Figure 7d-f.** Concurring with the earlier results relating to adaptation and sensitization, most ganglion cells are fed by two or more bipolar cell types and show combination of transient and sustained response features. **Figure S7a** shows the STA of a representative adapting and sensitizing RGC to blue stimulus. This is evident in the most STAs that consist of a fast phasic (~ 100 ms) and a slow integrative component (~ 1 s).

**Figure 7.**
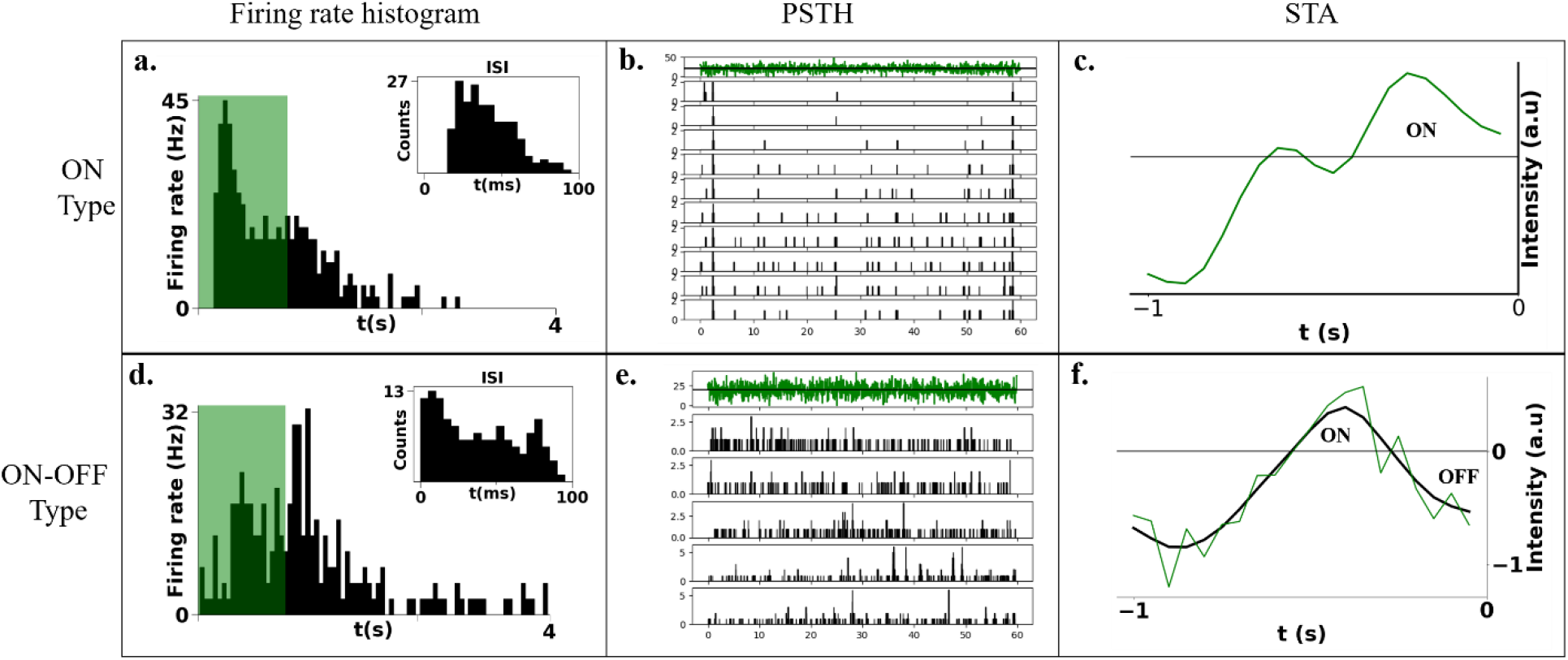
(a)The firing rate histogram of a representative ON RGC to green LED flashes(1s ON-5s OFF) across 10 trials. The inset shows the average ISI distribution. (b) PSTH of the same ON RGC to 10 trials of high (50 %) contrast Gaussian flicker sequence (20 Hz Frame rate, 60 s duration, identical across trials). (c) The corresponding spike triggered average (STA) of the ON RGC.(d)The firing rate histogram of a representative ON-OFF type RGC to periodic green LED flash. (e) PSTH of the corresponding RGC to Gaussian white noise flicker sequence. (f) The STA of the ON-OFF RGC.

### Precision of RGC firing events

While most RGCs were slow, a few examples of sparsely and precisely firing RGCs were observed.

The STA of a phasic OFF RGC is described in **Figure 8(a)**, showing that it responds to sustained low contrast dips of a 400 ms duration. The response of this RGC was back calculated from a full field gaussian random flicker stimulus with a variance that was 50% of the ambient luminance value. We observed three such RGCs in the same recording that responded to the same stimulus features; the average PSTH of the three OFF RGCs to ten trials of the same gaussian full field flicker stimulus is described in **figure 4(b)**. A raster plot of the firing event elicited after the stimulus feature is presented is described for one of the OFF cells during the ten trials of the repeated gaussian (**figure 4(c)**). We observed highly precise firing events in these RGC responses with a temporal jitter of the order of 10 ms. The corresponding firing rate histogram, PSTH for a repeated blue Gaussian flicker and temporal jitter for a representative blue ON RGC is described in **Figure S9** .The neonatal nature of the retina resulted in a significant amount of spontaneous activity which affected our binning/labeling of a firing event and resulted in the STA becoming noisy. (Hilgen et al., 2017)

**Figure 8.**
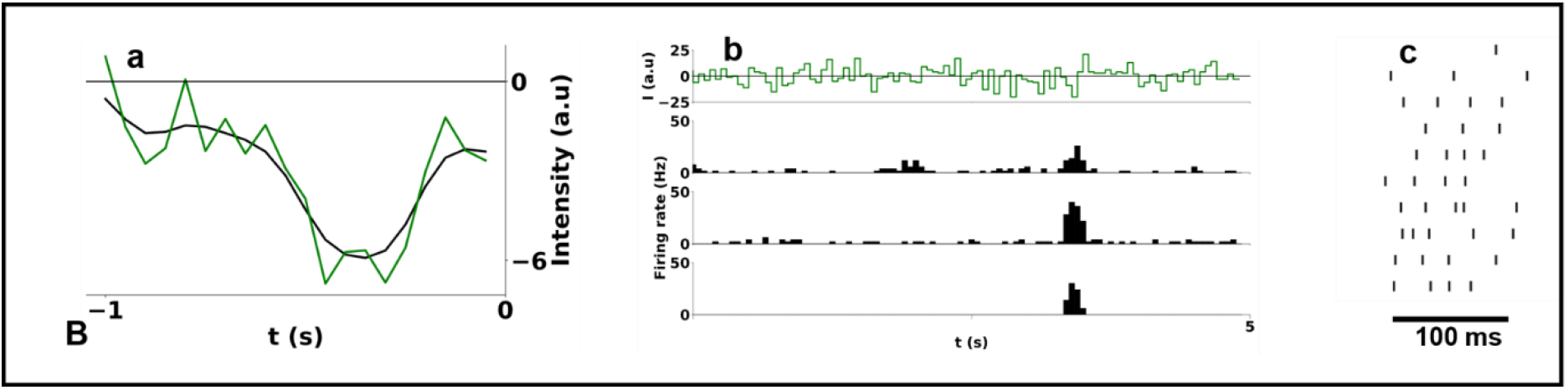
The STA for a representative OFF RGC is described in figure 4(a).4(b) The PSTH of 3 OFF RGCs to 10 trials of a 30 s gaussian full field flicker(50% contrast) stimuli. (c) The enlarged view of the firing events of OFF RGC is shown in a 100 ms scale. The neuron shows sparse and precise firing with a temporal jitter of ~10 ms.

### Color Dependence of Responses

The chick retina is a cone rich retina with a tri-stratified outer plexiform layer with rod and double cone axon segments terminating in the outermost strata, the green and red cones terminating in the middle segment and the blue and violet sensitive cones terminating in the innermost segment of the outer plexiform layer. The ability to discriminate between short (400-450 nm) and longer wavelengths (500-550 nm) is observed in all organisms that live in terrestrial habitats from honeybees to primates, (Osario and Vorobyev 2005) (Mills et al., 2014) and the cone rich avian retina is no exception to that. Color discrimination behavioral experiments done on chicks have implicated at least 3 different color opponent circuits at play. Electrophysiological evidence for their existence has not been shown so far to our knowledge. Despite the anatomical studies that describe the densely stratified plexiform layers (Seifert et al.2020) and a wealth of behavioral color discrimination studies in birds that implicate color opponency in the avian retina, electrophysiological studies that attest to this fact are lacking. The chick retina consists of 6 different types of photoreceptors. It has four single cone types expressing four different types of opsins namely, violet cones (419 nm) expressing a short wavelength sensitive 1 type opsin (SWS1), a short wavelength sensitive cone expressing a SWS2 type opsin absorbing maximally at 455nm (blue light), a medium wavelength sensitive cone expressing a rhodopsin-like 2 (RH2) with peak absorption at 508 nm corresponding to green light and long wavelength sensitive (LWS) opsin expressing cones with a peak absorption at 570nm corresponding to red light. A double cone and a rod photoreceptor that are touted to encode only luminance information are also expressed across the chick retina.

The effect of changing luminance of the full field flash varied across different putative RGC types observed and described reliable firing events at threshold intensities that depended on the wavelength of light. Increase in firing rate with increase in ambient illumination levels was only observed in clusters that showed slow sustained responses and was absent in clusters that showed fast and transient responses. Retinal explants from stages P0- P5 lacked a response to red light flashes in over 10 explants sampled across regions of the retina. This points to the likelihood that red responses develop only beyond the first week after hatching. We observe a blue OFF RGC, a violet OFF RGC and a green ON RGC that fire specifically to the wavelength and did not respond to other narrow band LEDs pointing to distinct spectral tuning of select RGC types. The sustained component of most responses was truncated or prolonged to different degrees depending on the wavelength of the stimulus. (**Figure S9, S10**) The firing event duration for sustained responses depended on the stimulus pulse duration and the duty cycle with the peak firing rate having the same approximate latency.

We first report the dominantly observed general trend with respect to blue and green light response of the putative RGCs. **Figure 9(a,b)** and **9(c,d)** shows the ON-response for green and blue stimuli, respectively. Importantly, Figure 9b and 9d shows contrasting luminance adaptation properties to the illumination wavelength. The green response shows a sharp adaptation, while the blue response shows an additional dominant sensitization feature. This is the dominantly observed feature in majority of the cell-types. Further, STA shown in the insets show contrasting response profiles corresponding to the respective cell-types. The positive peak in inset of **Figure 9b** corresponds to the fast adaptation feature, while the two regions indicated in the inset of **Figure 9d** corresponds to the adaptation and sensitization features.

**Figure 9.**
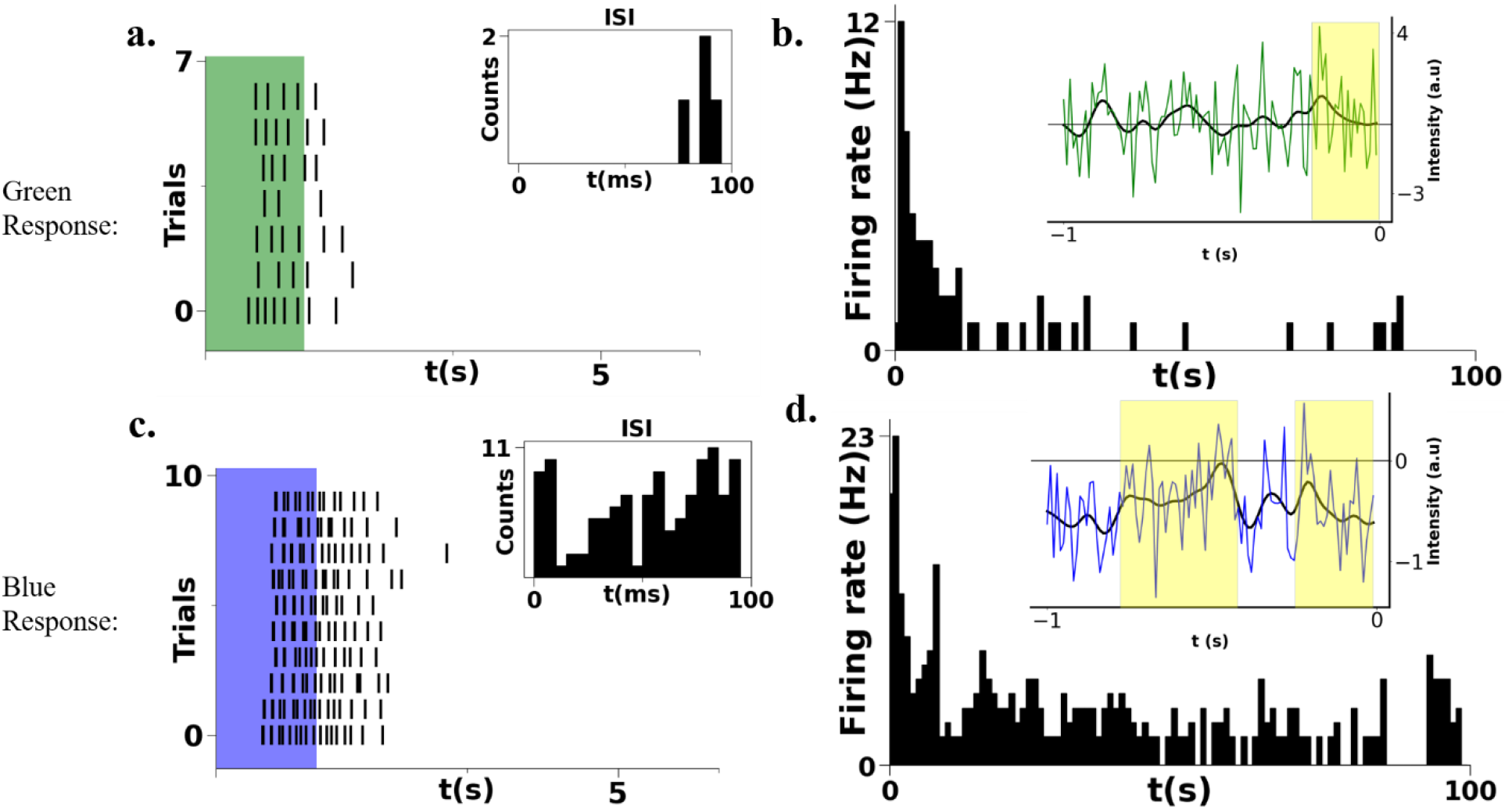
Color dependent response of the cells: (a)Raster plot representative adapting RGC to green light flashes. inset shows the average ISI distribution. (b) Firing rate histogram in same cell to uniform illumination. Inset: STA shows the ON response polarity prior to the spiking event. (c) Raster plot representative adapting and sensitizing RGC to blue light flashes. inset shows the average ISI distribution. (d) Firing rate histogram to uniform blue illumination. Inset: STA shows the ON and OFF response polarity prior to the spiking event.

The feature of color dependency was observed to be stronger in certain cell types that exhibit a green-ON/blue-OFF response. This “color-opponent” feature is illustrated in **Figure 10**. **Figure 10a and 10d** shows the opposing ON and OFF behavior to green and blue light pulse. This is in contrast to the “non color-opponent” cell type described earlier in **Figure 9**. **Figure 10b and 10e** describe the luminance adaptation behavior for green and blue light respectively. The STAs for green and blue are shown in **Figure 10c and 10f**, respectively. The STAs show contrasting polarity corresponding the peak firing latency of the full-field flash (shown in Figure 10a and 10d)

**Figure 10.**
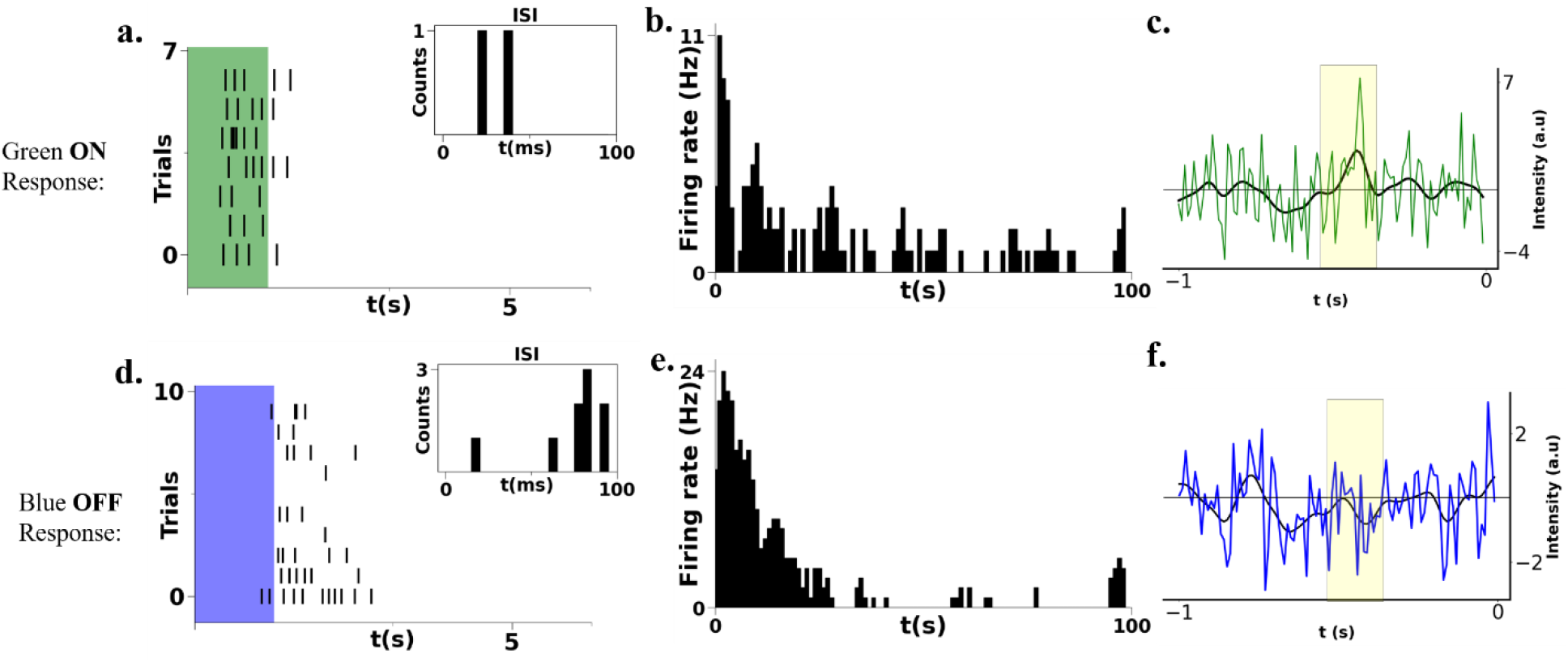
Color opponency in a representative RGC: (a)Raster plot representative ON-RGC to green light flashes. inset shows the average ISI distribution. (b) Firing rate histogram in same cell to uniform illumination shows adaptation and sensitization. (c) STA shows ON response polarity ~ 500 ms prior to the spiking event. (d) Raster plot representative OFF-RGC to blue light flashes. inset shows the average ISI distribution. (e) Firing rate histogram to uniform blue illumination shows dominant sensitization. (f) STA shows OFF-response polarity ~ 500 ms prior to the spiking event.

The color opponent responses can be attributed to the feedback inhibition from the horizontal cell to the cone. Yellow-blue (YBH) and red-green (RGH) c type horizontal cells have been implicated in the origin of color opponent responses in the turtle retina.(Twig et al. 2003).

The reptilian turtle retina expresses four different types of horizontal cells, just like in the avian retina. (Fischer et al., 2007) This might implicate the outer plexiform layer in color opponency in the avian retina just as in the primate, reptilian, and teleost fish retinae. (Crook et al., 2011) Yet another factor that leads us to believe that color dependency of the responses are horizontal cell-mediated is because they show a contrast level dependent switch in their polarity without compromising on their opponency (**Figure S11**). This hypothesis is further solidified from the outer plexiform layer connectivity study, where the double cone, which is believed to perform luminance-based coding, connects to different substrata of the outer plexiform layer differently with the principal member (L cone-bearing) and the accessory member (S cone bearing) describing distinct invaginating and basal contacts with other cones in the blue/violet and red/green sublayers. (Gunther et al., 2021)

Three types of correlations were observed due to the concerted firing between RGCs in the neonatal chick, namely a narrow correlation between neighboring sustained firing RGCs(Figure S), a medium correlation between neighboring transient firing RGCs and a broad correlation between RGCs as much as 1 mm apart.

**Figure 11** describes two sensitizing delayed ON fast OFF RGC that are recorded from adjacent electrodes in the MEA and their responses to green and blue light full field flash stimuli are shown in **Figure 11((a,b)&(d,e)**). As seen from their flash responses, the cells fire very similarly to green and blue light stimuli, but as shown in **figure 11(c&f)** their cross-correlation functions are very different. Blue light stimuli elicit higher firing rates but green light stimuli result in much higher correlation strengths, indicative of color information being encoded in pairwise correlations even when the nature of firing is similar. Thus, color information encoding in the chick retina through correlations between adjacent cell types allows for color information transfer beyond the conventional color opponency seen as opposing contrast sensitivity to light flashes of different colors. (Zhou et al., 2005)

**Figure 11.**
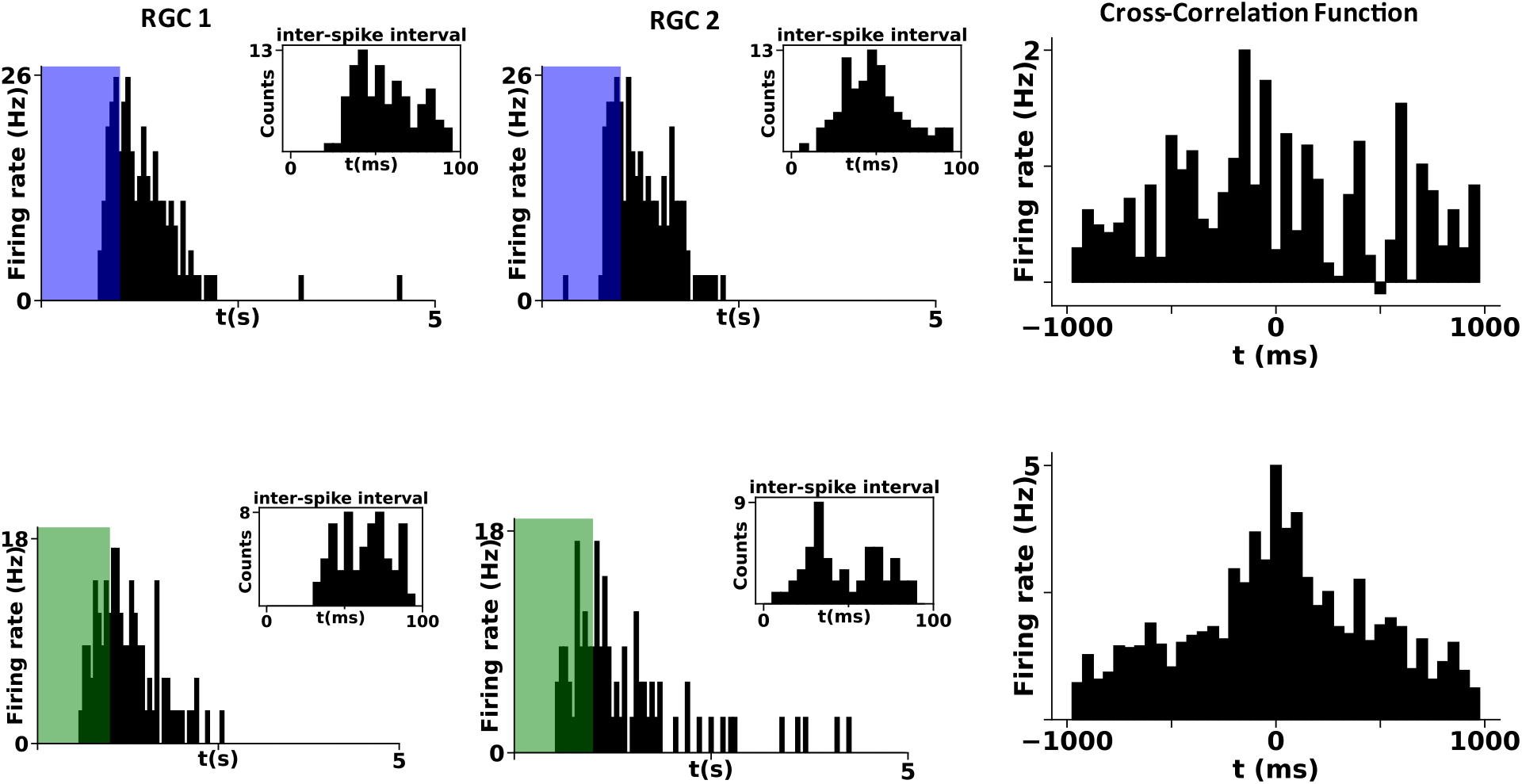
Color dependent cross-correlation: Raster plots of neighboring sensitizing RGCs to green light flashes for (a) RGC 1 and (b) RGC 2. (c) Corresponding CCF shows strong broad correlation for a green light stimulus. Raster plots of same neighboring sensitizing RGCs to blue light flashes for (d) RGC 1 and (e) RGC 2. (f) Corresponding CCF shows weaker correlation for a blue light stimulus.

## Discussion

Extracellular recording of chick retinal explants from postnatal p0 to p5 yielded clean single units that described rich distinctive features that depended on luminance contrast color and the temporal frequency of the stimulus. The elicited firing across luminance levels and fixed contrast steps of different colors indicated features of both rate coding and precision based temporal coding. Some RGCs showed robust firing to slow periodic flashes (1 s ON- 5 s OFF) and did not respond to rapidly flickering stimuli (10-50 ms refresh rate), while others showed clean reliable responses to random flicker stimuli but not to the periodic flash. This points to integration time differences between the center and surround of the receptive field which is used to compute spatial and temporal characteristics of the stimulus.

Single units were sorted clustered based on threshold crossings and clean autocorrelation functions. The polarity and kinetics of the RGC type can be defined by the nature of contrast changes that elicited a firing event i.e. its spike-triggered average but as the neonatal retina still shows a significant degree of spontaneous activity and as the firing events are protracted, the STAs for most isolated single units were noisy. So we corroborate the STAs with RGC responses to other stimuli such as ambient illumination and full field contrast steps. It has been observed that bipolar cell synapses show sensitizing or adapting features depending on stimulus features such as frequency, luminance, contrast. (Niloleav et al., 2013) The present studies show that the sensitizing and or adaptive features of ganglion cells also depends on the color and irradiance of the stimulus. The results also indicate that blue light stimulus resulted in sharp sensitization only responses or sharp adaptation and sensitization features to constant luminance, while very few RGCs showed contrast adaptation to blue light. Green light responses on the other hand mostly showed luminance and contrast adaptation with a few responses showing an additional broad sensitization feature. The RGCs presumably respond to inputs from two types of bipolar cell types as they showed both the adaptation and sensitization features. The tonic and phasic bipolar cell inputs feed sustained and transient RGC types respectively. (Awatarmani and Slaughter, 2000) A large fraction RGC responses were of a mixed type (tonic and phasic) pointing to both bipolar cell types synapsing onto RGCs with horizontal cell input influencing the depressing or facilitating nature of the connection in a color dependent manner. (Twig et al., 2001) Some RGCs in the neonatal chick retina showed ambipolar STAs arising likely due to fundamental nonlinearity/ multimodality in its response. This would require further spike-triggered covariance (STC) to extract the temporal receptive field profile. Most of the RGCs recorded from the peripheral Chick retina showed very low contrast sensitivity, with reliable firing elicited only for very high contrast changes (~100%). This can be attributed to the high density of retinal ganglion cells in the chick retina, which thrives in a wide range of luminance conditions. (Baden et al., 2020) The high visual acuity of the chick retina in the dorsal area and the area centralis could mean a very high contrast threshold is feasible to detect wavelength-dependent contrast changes a spatially dense receptive field mosaic. This could mean the chick compensates for low contrast sensitivity with high visual acuity (high spatial contrast sensitivity). Color opponency was evident in the polarity of the contrast values that elicited activity in different temporal windows before a firing event for gaussian stimuli of different colors. A behavioral study monitoring the object fixation in chicks compared to mammals showed that chicks investigate objects by looking at them with different specialized parts of the eye, unlike primates that fixate objects onto the fovea. (Stamp Dawkins, 2001) The presence of centrifugal projections in the area centralis of the chick retina could enable the utilization of specialized regions in the retina to observe different aspects of the visual scene and convey complementary sets of information about it. This is also in sync with different regions having different wavelength-dependent contrast sensitivities. It was observed that even a small difference in location of the retinal explant piece in the dorsal area resulted in a large difference in the population of cell types that were obtained. The morphological similarity of the four horizontal cell types in the chick retina to the horizontal cell types found in the turtle retina has been shown previously.(Fisher et al., 2006) Thus this color opponency is likely mediated by a yellow-blue horizontal (YBH) cell as it has been previously reported in the turtle retina.(Twig et al., 2002) These color opponent cells show opposing contrast sensitivities observed in the time course of the STA at ~500 ms before the onset of spiking. Apart from these conventional color opponent cells, color information is also likely encoded in other features of many non-color opponent cells such as the latency, nature of the response or population coded in the cross-correlation strength for many neighboring RGC pairs (Chen et al., 2005) We record from a large subset of slow RGCs in the dorsal area of the chick retina that show a combination of differential adaptation and sensitization features that modulate their temporal tuning curves Thus, chick RGCs behave as complex temporal coders that respond to wavelength-dependent contrast values at specific temporal windows before an elicited firing event. These observations are indicative of the diversity of cell types and complex interconnectivity in the outer plexiform layer of the avian retina. This architecture enables and contributes to the processing of the visual scene into several behaviorally relevant features. Thus the chick retina describing rich distinctive features in RGC firing patterns to different stimuli can be used has a good yardstick for the validation of subretinal prosthetic devices.

## Methods

### Ethics statement

All studies were performed in accordance with the guidelines of the Institutional Animal Ethics Committee (IAEC) and the Institutional Bioethics and Bio-safety Committee, Jawaharlal Nehru Centre for Advanced Scientific Research (JNCASR). All protocols and experiments were approved by the IAEC and were performed in accordance with the guidelines followed by the Poultry Science Unit, KVAFSU (Karnataka Veterinary, Animal and Fisheries Sciences University, Hebbal, Bangalore)

### Animal Preparation and Retina isolation

Fertilized Giriraja Gallus gallus domesticus eggs were procured from the Poultry Science Unit, KVAFSU and incubated in a Brinsea Octagon 28 incubator till they reached the appropriate stage of development (P0-P5). Neonatal chicks were obtained on the day of the experiment in a well-ventilated, well-lit portable brooder. The neonatal chicken was sacrificed through cervical dislocation. The eyes were enucleated, the eye cup was hemisected, and then the retina was isolated in ice-cold aCSF under dim light conditions. The dissected retinae were preserved in freshly prepared aCSF solution with 124 mM NaCl, 25 mM NaHCO_3_, 1.2 mM HEPES, 5 mM KCl, 1.2 mM MgSO_4_, 2.5 mM CaCl_2_, and 10 mM glucose at pH 7.4 that was constantly bubbled with carbogen gas (95% CO_2_ and 5% O_2_). A suitable healthy retina was then removed to a separate bubbled petri dish under a stereo microscope, where it was cut to the desired size and placed on an annularly cut custom nylon mesh with a pore size of 100 μm with the photoreceptor side facing the nylon mesh. (Deepak et al. 2022) There are intrinsic difficulties of recording from the chick retina owing to waves of spreading depression after the onset of which the fundamental firing properties of RGCs change and the native response state is nearly impossible to recover. A popular strategy that is employed to prolong the survivability of the explant is the application of the Mg^2+^ block by way of increased MgCl_2_ in the recording medium and its wash-off prior to the electrophysiological recording. In the absence of clear data on the time course of the wash-off of the Mg^2+^ block in chick retinal explants we chose to probe the light response of the chick retinal explants in carbogen gas bubbled normal chick ACSF in the absence of the Mg^2+^ block. Thus, only samples that last over 45 minutes before the onset of the spreading depression event are considered for the analysis.

### Spike Sorting and Analysis

Spyking-Circus (Yger et al., 2018) was used to spike sort the retinal recordings. In the first step, spikes from the band-pass filtered data (300 – 3000 Hz) are detected as threshold crossing (7 S.D) and snippets of data around each crossing of appropriate duration (Nt = 5 ms) are cutout across all electrodes (Ne = 59). A maximum of Ns = 10000 snippets are gathered from each electrode for a total of Ns *Ne waveforms. The waveforms are in a high dimensional space due to the high sampling rate. Principal Component Analysis (PCA) was done, to reduce the dimension of the problem. This number was taken to five by setting the parameter NPCA = 5. The clusters thus obtained from spyking circus were then subjected to post-processing which involved cleaning of the templates to ensure that the percentage of refractory period violations per cluster was under 1%. Templates that showed higher than 1% refractory period violations and with spike counts under 500 were discarded from the analysis.

**Firing rate** was calculated by aligning spikes to the onset of the light stimulus for ten trials. The histogram is the average spikes in each 50 ms bin.

**Cross-Correlogram** between two putative ganglion cells was calculated by aligning the spikes of the second neuron to every spike of the first neuron and the average number of spikes of the second neuron was counted in 10ms bins. To remove stimulus correlations from the cross correlograms a **shifted cross-correlogram** was calculated by shuffling the trials and computing the cross-correlogram. This was subtracted from the initial cross-correlogram to obtain the **shift corrected cross-correlogram.**

**Spike Triggered Average (STA)** was calculated by taking the stimulus values till one(two) second(s) before each spike and averaging them during the Gaussian White Noise stimulus for all trials. The smoothened STA trace was obtained by convolving the STA with a Gaussian kernel.

The Gaussian White Noise (GWN) stimulus was shown by changing the intensity value of the LED with a refresh rate of 10ms (25/50) picked from a Gaussian distribution. The contrast of the GWN stimulus was defined as dI/I0 where dI is the variance of the Gaussian distribution and I0 the mean intensity

## Supporting information

Supplementary Info neonatal chick retina

## Acknowledgments

We acknowledge Dr. N Ganesh for his valuable suggestions and inputs with regards to the presentation of different key findings and results. We also thank him for engaging in long drawn-out discussions that led to many of the ideas/ inferences presented here. We also acknowledge late Anil Krishna Konduri for his contribution and assistance in improving recording quality and artifact suppression in the initial phase of the project.

