## Supplementary Info neonatal chick retina for "Temporal Characteristics of Neonatal Chick Retinal Ganglion Cell Responses: Effects of Luminance, Contrast, and Color"

### Supplementary Information

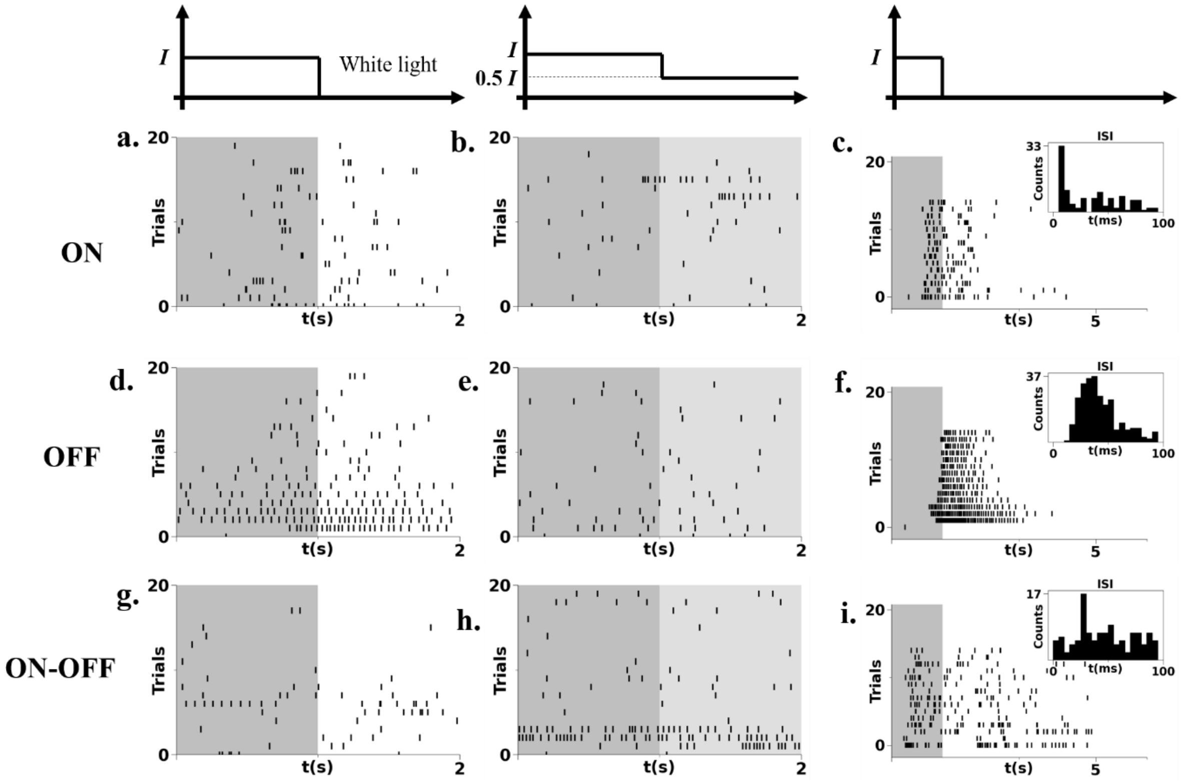


Figure S1: responses of representative RGC's of different polarities to different white light full field flash protocols. The responses of a representative (a) ON, (d)OFF and (g)ON-OFF cluster to white light flashes from the dark (1 s ON- 1 s OFF). (b,e,h)The response of the same ON, OFF and ON-OFF RGCs to 100% full field contrast steps from an ambient luminance of 7 $mW/m^2$ at a  duty cycle of 50%. (c,f,i) The response of the same representative ON,OFF,ON-OFF RGCs to a 1s ON-5s OFF white light from the dark.

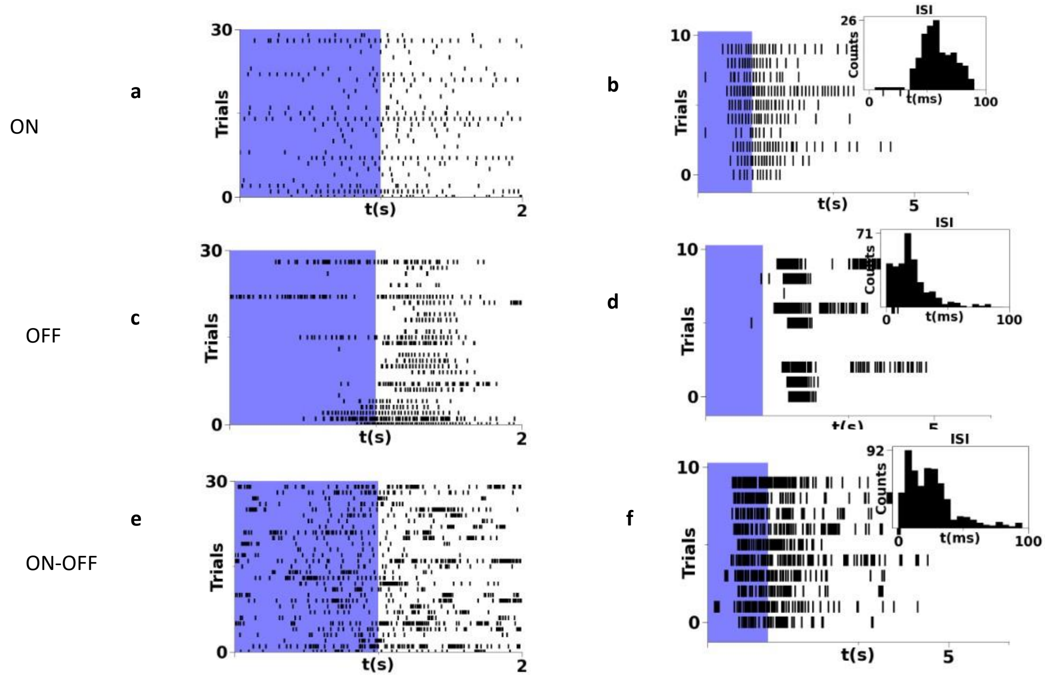


Figure S2: (a)The response of a representative delayed ON RGC to blue full field flashes (50% duty cycle,1mW/$m^2$) from the dark. (b) The response of the same ON RGC to 20 % duty cycle blue light flashes at the same intensity. (c,d)The responses of a representative OFF RGC to 50% and 20% duty cycle flashes respectively.(e,f) The responses of an ON -OFF RGC to 50% and 20% duty cycle blue light flashes from the dark.

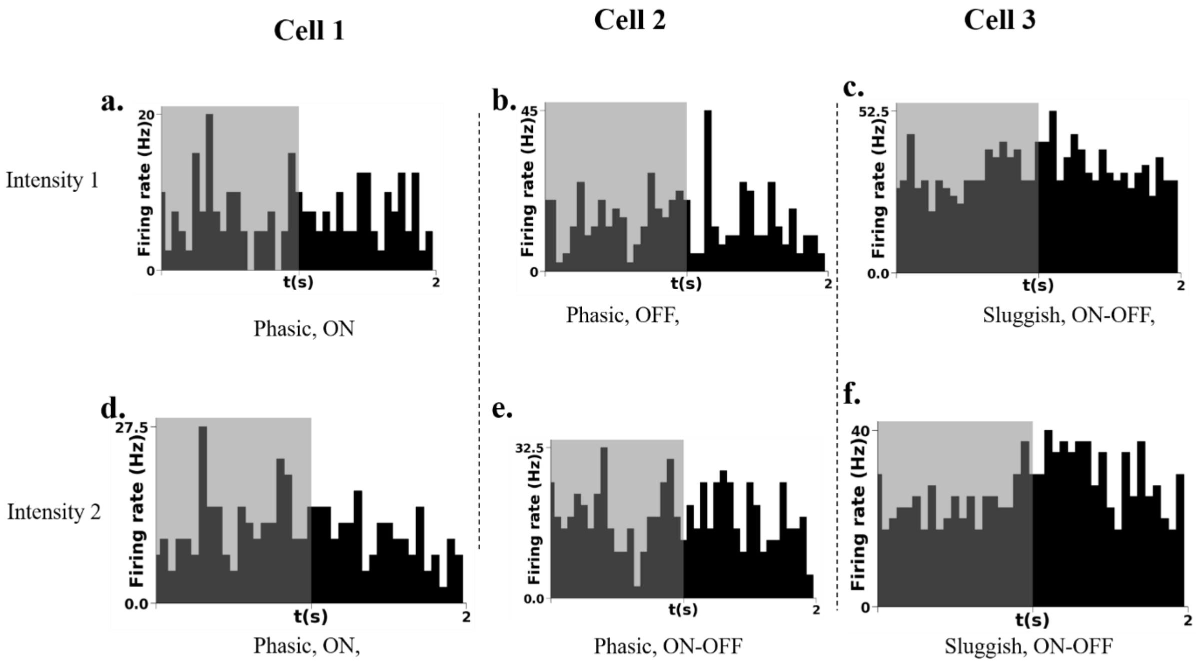


Figure S3: the response of representative on off and on off RGC's across two different luminance levels. A phasic response of **cell 1** at intensity of 14 mW/m2; b a representative phasic off response to fulfill white light stimulus at 50% duty cycle see the response off by representative on off cell do the same stimulus protocol. (d) the response of **cell 1** add an intensity of 35 mW/m2 to a full field white light flash at 50% duty cycle (e) the response of **cell 2** at an intensity of 35 mW/m2 resulting in a polarity switch to a ON-OFF type response (f) response of **cell 3** do the same stimulus at the 35 mW/m2 intensity.

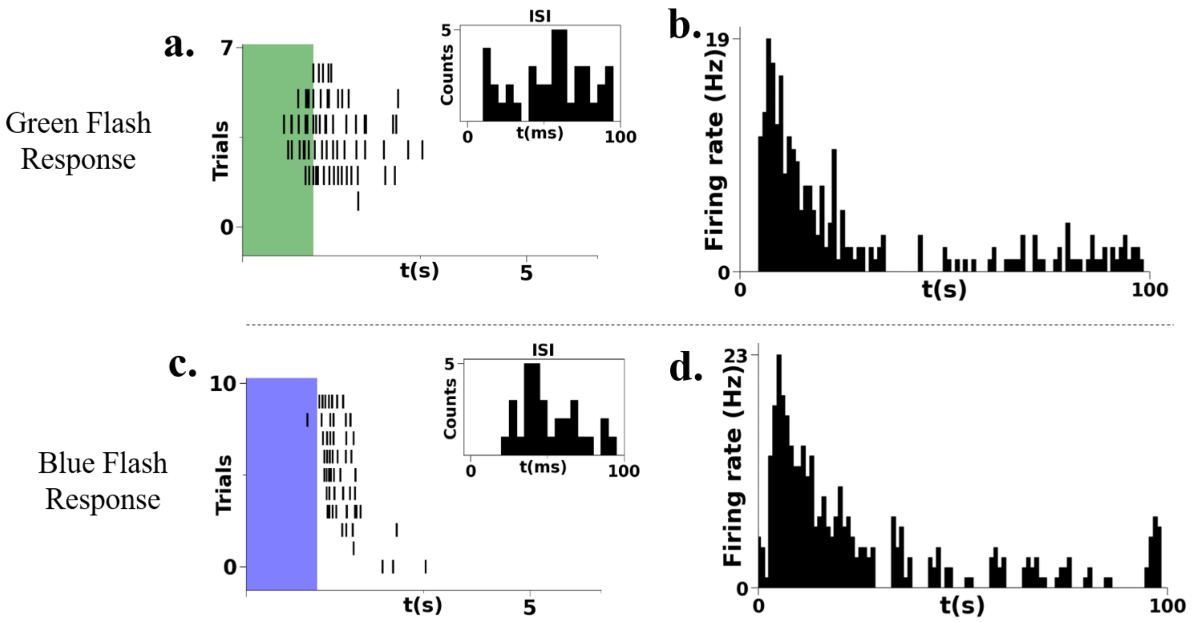


Figure S4: (a)The raster plot offer sensitizing off ganglion cell to green full field flash over 10 trials (b) the firing rate histogram for the same RGC to constant green light illumination (c) the raster plot off the same sensitizing ganglion cell to blue light flashes (d) the firing rate histogram for constant blue light illumination showing a clear sensitizing feature

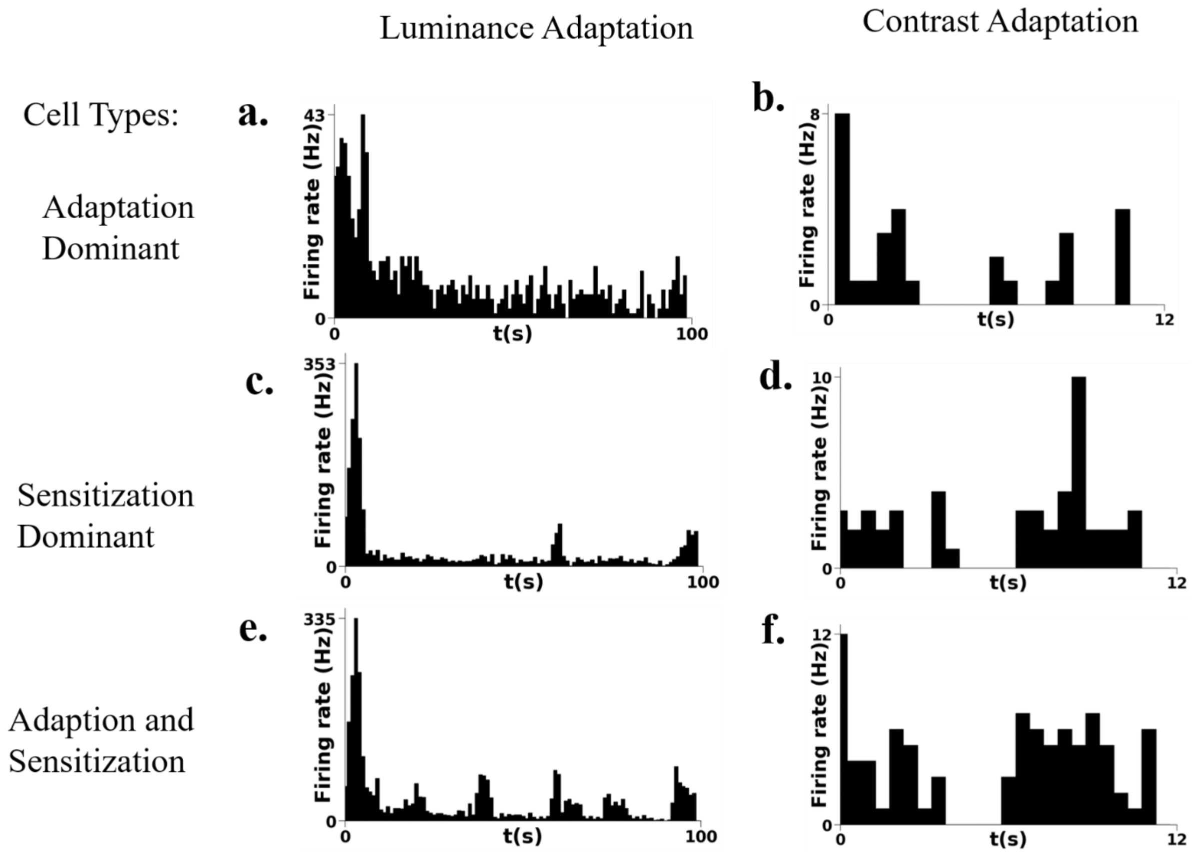


Figure S5: adaptation and sensitization to blue light (a) The response of an adaptation dominant RGC to ambient illumination with blue light (b) describes a slow contrast adaptation feature when presented with a high contrast Gaussian stimulus sequence (50%, refresh rate 10 ms, 12 s long) following a low contrast sequence (25%); note that the response is averaged across different high contrast gaussian sequences. (c) The response of a sensitization dominant RGC to ambient illumination; (d) the averaged RGC response upon moving to a high contrast gaussian stimulus (e) the firing rate histogram of an RGC to ambient illumination showing only the sensitization feature while (f) describes the response of the same RGC when presented with a high contrast Gaussian stimulus showing features of both slow contrast adaptation and sensitization. Thus, Blue light shows contrasting features with respect to luminance and contrast.

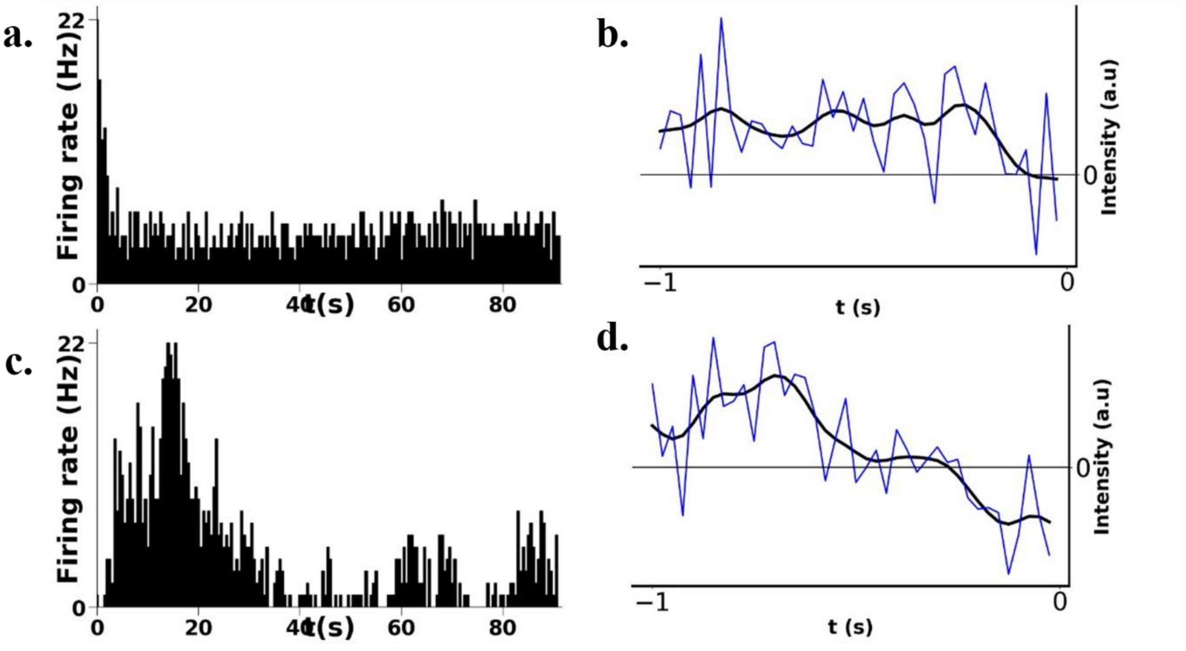


Figure S6: (a)The response of a fast OFF adapting RGC to constant blue light illumination (b) STA of that same RGC calculated from gaussian white noise stimulus sequence showing a slow integrating on response followed by phasic off response (c) the response of a sensitizing off RGC to constant blue light illumination (d) the STA of that same ganglion cell calculated from a gaussian white noise sequence showing a  slow  OFF component of the order of 200 ms.

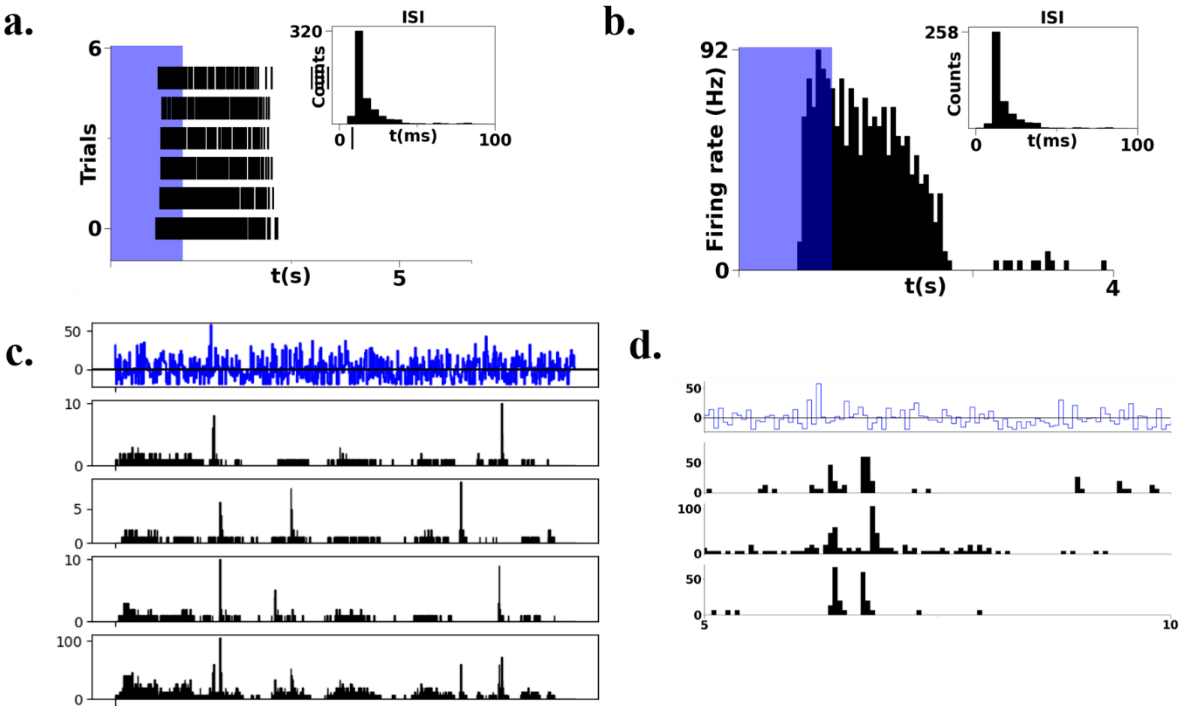


Figure S7 (a)The raster plot of the response of an ON RGC to blue light flashes (6 trials) (b)The firing rate histogram of the same cell (c)The PSTH of the same ON RGC to 4 repeated trials of a random Gaussian white noise sequence(20 Hz, 1200 frames) (d) The response of three different ON RGCs of the same type averaged across the 4 trials of the random gaussian sequence showing low jitter and high precision

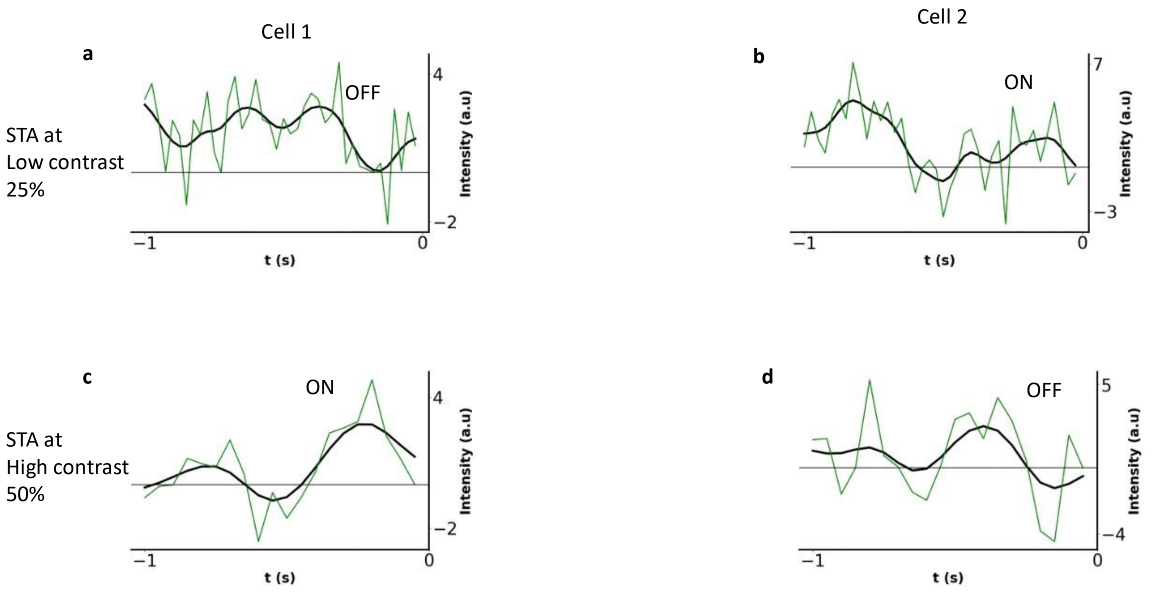


S8 (a) The STA of a putative RGC (cell 1) to green light at a mean light intensity of 3.4 $mW/m^2$ at 25% contast,(c) The STA of the same RGC when presented with a contrast of 50% (b,d) The STA of a putative RGC to low(25%) and high contrast Gaussian flicker respectively

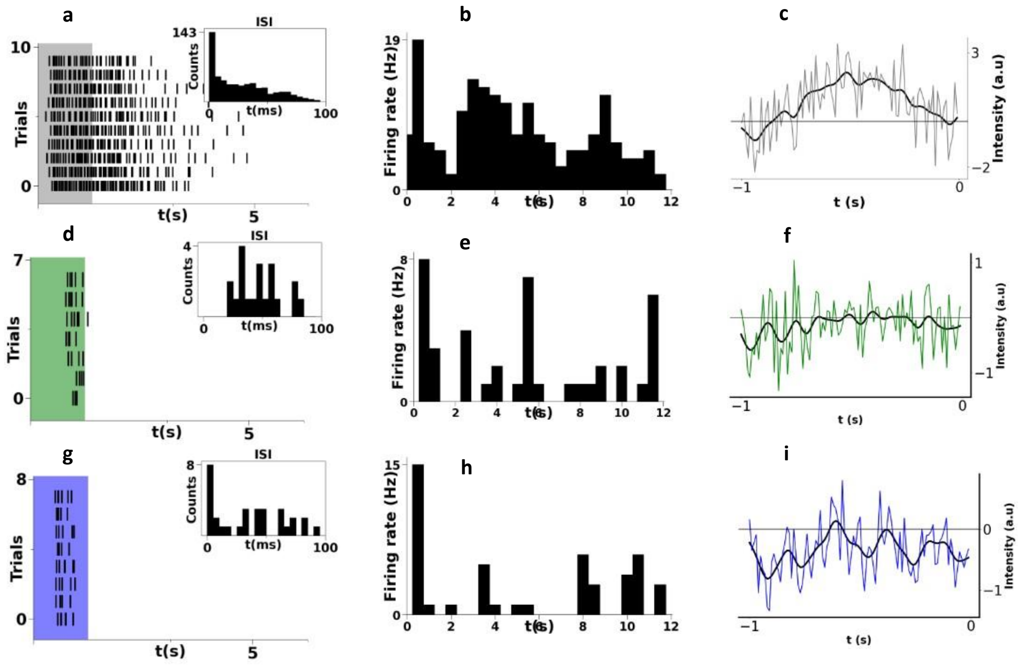


Figure S9 Nature of response varying with color (a)Full field flash responses of a cluster to full field white light flashes 1s ON-5 s OFF duration. (b)The firing rate histogram of the same cluster to a high contrast (50% from 25%) random Gaussian flicker sequence averaged across 10 random Gaussian sequences describing slow contrast adaptation and sensitization at two different time scales. (c)The STA of the RGC to white light stimuli. (d) The full field flash response of the same RGC cluster to green light flashes (20% duty cycle) (e)The firing rate histogram of the same cluster to high contrast Gaussian flicker sequence averaged across 10 random Gaussian sequences (f) The STA of the same cell to green light stimuli (g) The full field flash response of the RGC to flue light flashes (20% duty cycle) (h) The firing rate histogram in response to a high contrast(50% from 25%) random gaussian sequence (i) The STA of the RGC to blue stimuli

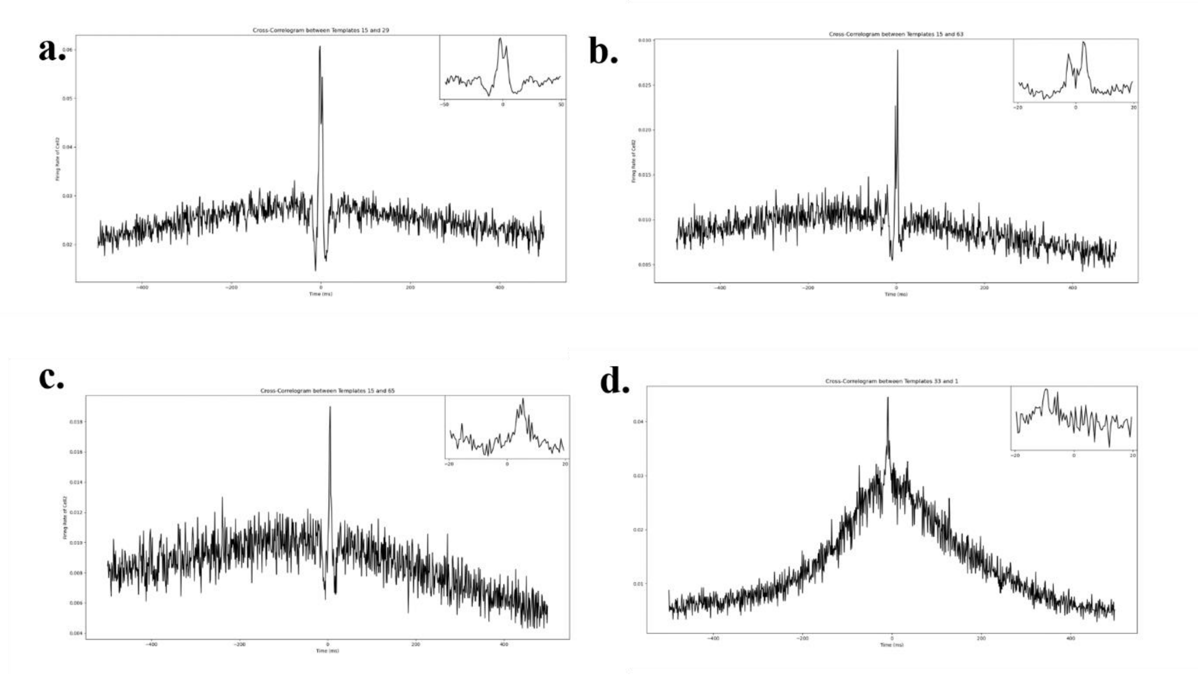


Figure S10: The three basic types of correlations observed between neighbouring RGCs in the chick retina. (a ,b) Gap junction mediated narrow correlations between two neighbouring ganglion cells with variable reciprocal coupling between the two cells. (c)A medium correlation between two ganglion cells which is likely mediated by an inhibitory amacrine cell. (d) A broad cross correlation between neighbouring ganglion cells

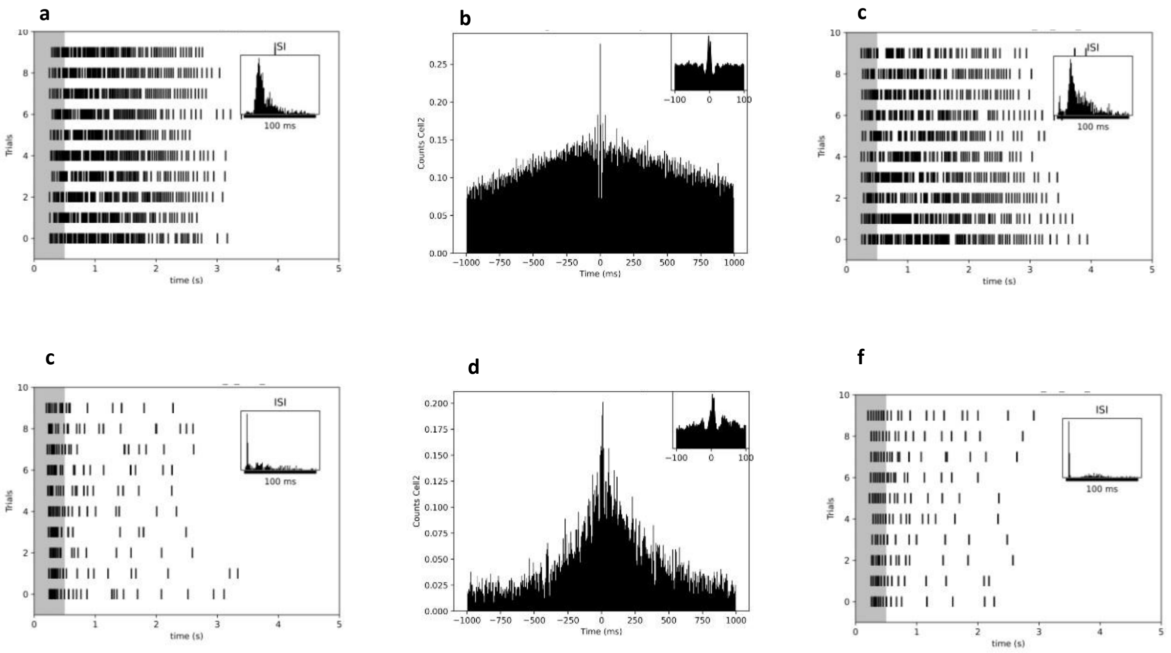


Figure S11: (a,c)The raster plots of two sustained ganglion cell types to 10 trials of a 500 ms full field flash at a duty cycle of 10%. (b) The cross-correlation function computed between the two adjacent sustained firing RGCs describing a narrow correlation. (d, f) The raster plots of two

neighbouring transient firing RGCs to 500 ms full field flash (10% duty cycle) and (e) their cross-correlogram describing a medium type of correlation.
